# The SAGA core module is critical during *Drosophila* oogenesis and is broadly recruited to promoters

**DOI:** 10.1101/2021.06.18.448946

**Authors:** Jelly H.M. Soffers, Sergio Garcia-Moreno Alcantara, Xuanying Li, Wanqing Shao, Christopher W. Seidel, Hua Li, Julia Zeitlinger, Susan M. Abmayr, Jerry L. Workman

**Affiliations:** Stowers Institute for Medical Research, Kansas City, Missouri, 64110, USA; Department of Pathology and Laboratory Medicine, University of Kansas School of Medicine, Kansas City, Kansas, 66160, USA; Department of Anatomy and Cell Biology, University of Kansas School of Medicine, Kansas City, Kansas, 66160, USA

## Abstract

The Spt/Ada-Gcn5 Acetyltransferase (SAGA) coactivator complex has multiple modules with different enzymatic and non-enzymatic functions. How each module contributes to gene activation in specific biological contexts is not well understood. Here we analyzed the role of the non-enzymatic core module during *Drosophila* oogenesis. We show that depletion of several SAGA-specific subunits belonging to the core module blocked egg chamber development during mid-oogenesis stages, resulting in stronger phenotypes than those obtained after depletion of SAGA’s histone acetyltransferase module or deubiquitination module. These results, as well as additional genetic analyses pointing to an interaction with TBP, suggested a differential role of SAGA modules at different promoter types. However, SAGA subunits co-occupied all promoter types of active genes in ChIP-seq and ChIP-nexus experiments. Thus, the SAGA complex appears to occupy promoters in its entirety, consistent with the strong biochemical integrity of the complex. The high-resolution genomic binding profiles are congruent with SAGA recruitment by activators upstream of the start site, and retention on chromatin by interactions with modified histones downstream of the start site. The stronger genetic requirement of the core module during oogenesis may therefore be explained through its interaction with TBP or its role in recruiting the enzymatic modules to the promoter. We propose the handyman principle, which posits that a distinct genetic requirement for specific components may conceal the fact that the entire complex is physically present.

**Author Summary:** Embryonic development critically relies on the differential expression of genes in different tissues. This involves the dynamic interplay between DNA, sequence-specific transcription factors, coactivators and chromatin remodelers which guide the transcription machinery to the appropriate promoters for productive transcription. To understand how this happens at the molecular level, we need to understand when and how coactivator complexes such as SAGA function. SAGA consists of multiple modules with well characterized enzymatic functions. This study shows that the non-enzymatic core module of SAGA is required for *Drosophila* oogenesis, while the enzymatic functions are largely dispensable. Despite this differential requirement, SAGA subunits appear to be broadly recruited to all promoter types, consistent with the biochemical integrity of the complex. These results suggest that genetic requirements and physical organization do not always go hand in hand.

## Introduction

The Spt/Ada-Gcn5 Acetyltransferase (SAGA) complex is required for the transcription of most RNA polymerase II (Pol II) genes [1]. It contains several multi-protein modules that are functionally distinct and have specialized functions in transcription. The histone acetyltransferase (HAT) module binds histone (H) 3 lysine (K) di- and trimethylated histones and preferentially acetylates H3K9 and H3K14, which leads to gene activation [2-5]. The deubiquitinase (DUB) module removes H2BK123ub, which directly affects transcription elongation and indirectly affects other histone modifications [6, 7]. The other two modules have no described enzymatic function. The module formed by the large Tra1 subunit (*Drosophila* Nipped-A) accommodates interactions with transcription activator proteins, which are important for promoter recruitment [8-13, 14, 15-17]. The core module is composed of Spt3, Spt20, Ada1 and several TBP-associated factor (TAF) subunits. Some of the TAFs are shared with the promoter recognition complex TFIID, and some functions such as TATA Binding Protein (TBP) binding overlap between SAGA and TFIID [18-20]. TBP binds DNA in a sequence-specific manner and helps to position the preinitiation complex. The structural basis for TATA Binding Protein (TBP) binding is highly conserved between the SAGA complexes across species and depends on interactions with Spt3 [21-23].

The precise non-enzymatic functions of the SAGA complex in promoter regulation have remained incompletely understood. Specifically, it has remained unclear if different promoter types rely on enzymatic versus non-enzymatic functions for their activation. An attractive genetic model system to study the role of the SAGA core module in promoter function is *Drosophila* oogenesis. In this model, the genetic requirement of the enzymatic SAGA modules has already been investigated, and it appears that while the activity of the SAGA DUB module is dispensable, the histone acetylation activity mediated by Gcn5 and Ada2b is critical for the progression of oogenesis [24, 25]. Though these results could be explained by a critical requirement for the Ada2b- and Gcn5 containing ADA or Chiffon complexes [26, 27] and no dependency on the SAGA complex during oogenesis, they may also reflect that distinct gene sets, defined by promoter types, have a different genetic requirement for each module. During *Drosophila* spermatogenesis and oogenesis, both TBP and the structurally related protein TBP-related factor 2 (TRF2) play critical, yet different roles, suggesting that different TBP-binding and/ or containing complexes may be required for oogenesis [28-39]. Therefore, we hypothesized that the SAGA complex itself and specifically the TBP-binding core module is important for oogenesis.

We investigated the role of the SAGA core module during *Drosophila* oogenesis by specifically targeting subunits of the *Drosophila* SAGA core complex that are not shared with TFIID: SAF6 [40], TAF10b [40, 41] and WDA [42]. We found that depletion of these SAGA-specific TAFs arrested egg chamber development during mid-oogenesis, demonstrating a critical role for the SAGA core module. Depletion of the TBP-binding core subunit Spt3 phenocopied this oogenesis defect, but the direct depletion of TBP or TFIID subunits resulted in a more severe phenotype, indicating that the latter play a broader role. The genome-wide occupancy of SAGA subunits, as observed in ChIP-seq experiments or in the higher resolution ChIP-nexus experiments [43], showed that SAGA subunits co-occupied all promoter types and correlated with transcriptional activity, but did not indicate that the SAGA complex specifically associates with TBP or TRF2 at distinct promoter types. However, at high resolution, differences in binding patterns between promoter types became apparent. The complex preferentially binds either upstream of the start site, where activators are found, or downstream of the start site, where SAGA may be retained at the +1 nucleosome.

Our results carry an inherent caution when interpreting genetic, phenotypic, and genomic data, since the distinct genetic requirement for specific modules may conceal the fact that the entire complex is physically present. During oogenesis, the non enzymatic core module functions are most critical, but this cannot be explained by the specific recruitment to distinct promoter types. We propose the handyman principle: while a handyman has many tools, he will hardly use them all at the same time, and which tool he uses will depend on what requires maintenance.

## Results

### SAGA modules have distinct roles during Drosophila oogenesis

Oogenesis in *Drosophila* occurs in an assembly line fashion in units called ovarioles, which are strings of six or seven sequentially developing egg chambers. The development of the egg chambers is divided into early stages (1-6), mid (stage 7-10a) and late stages (10b-14) [44]. During mid-oogenesis, the egg chambers take on an elongated shape and the oocyte increases in size.

We reported previously that the enzymatic components of the SAGA complex are differentially required for oogenesis [25]. The DUB module is dispensable for oogenesis since mature eggs form after the depletion of the enzymatic deubiquitinase subunit Nonstop in the germ line (Fig 1A). Ovaries lacking Ada2b (Fig 1B), the subunit that modulates Gcn5 acetyltransferase function, display defects in late oogenesis stages. These *ada2b[1]* germ line clone ovaries still have enlarged oocytes typical of late stages, but the dorsal filaments fail to form and the oocytes degenerate before they develop into mature eggs (Fig 1B).

**Fig 1.**
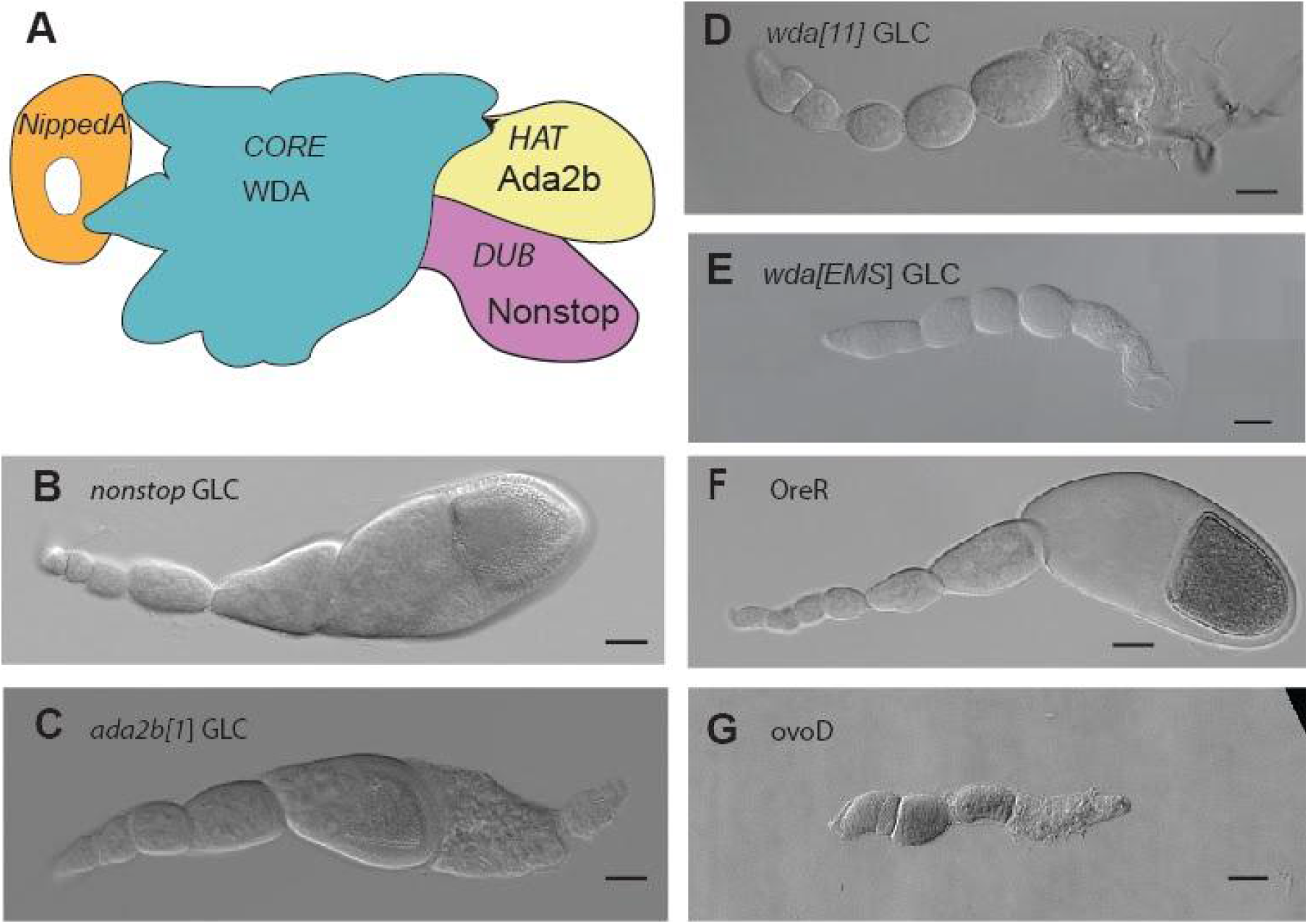
Different requirements of the SAGA modules during oogenesis. (A) Cartoon of the SAGA complex. (B-G) Differential contrast images of GLC ovarioles and controls. Loss of the enzymatic functions leads to no or a late defect in oogenesis, because DUB subunit Nonstop is dispensable, whereas the *ada2b[1]* GLC ovaries display a stage 12 defect. (B) *Nonstop* GLC ovariole. (C) *Ada2b[1]* GLC ovariole. (D) *wda[11]* ovariole. (E) *wda[EMS]* ovariole. Scale bar: 50 µM

To test the genetic requirement of the core module, we depleted the SAGA-specific core subunit WDA by generating germ line clones of *wda[11]* (Fig 1C). Since the *wda[11]* allele also disrupts a neighboring gene [42], we performed an ethyl methanesulfonate (EMS) mutagenesis screen to isolate a new mutant allele of *wda* (*wda[EMS])* (Fig 1D). In ovaries with *wda[11]* or *wda[EMS]* germ line clones, the number of round early-stage egg chambers increased and late-stage egg chambers did not form, indicating a failure to progress through stage 7 of mid-oogenesis. This represents a phenotype that is even more severe than that associated with the loss of the enzymatic HAT or DUB activities and thus suggests an important and early function for the SAGA core module during oogenesis.

### SAGA core subunits are critical for mid-oogenesis

To corroborate the role of the core module during oogenesis, we depleted other SAGA-specific core subunits in the germ line and asked whether this phenocopies the WDA oogenesis defect. We created a *saf6* deletion allele by replacing the coding region with a dsRED cassette using CRISPR Cas9 (see methods) and generated germ line clones with this allele (Fig 2A). Loss of SAF6 led to a phenotype similar to the loss of WDA (Fig 1D, E). The egg chambers failed to elongate but remained round, and the oocyte did not increase in size, suggesting that oogenesis arrested at stage 6. A similar phenotype was observed when SAGA-specific TAF subunits were targeted in the germ line by expressing UAS RNAi with the maternal triple Gal4 driver. The depletion of SAF6, WDA or TAF10b all arrested oogenesis at stage 6-7 with similar phenotypes (Fig 2B-D), demonstrating that SAGA-specific TAF subunits are critical for the progression through mid-oogenesis.

**Fig 2.**
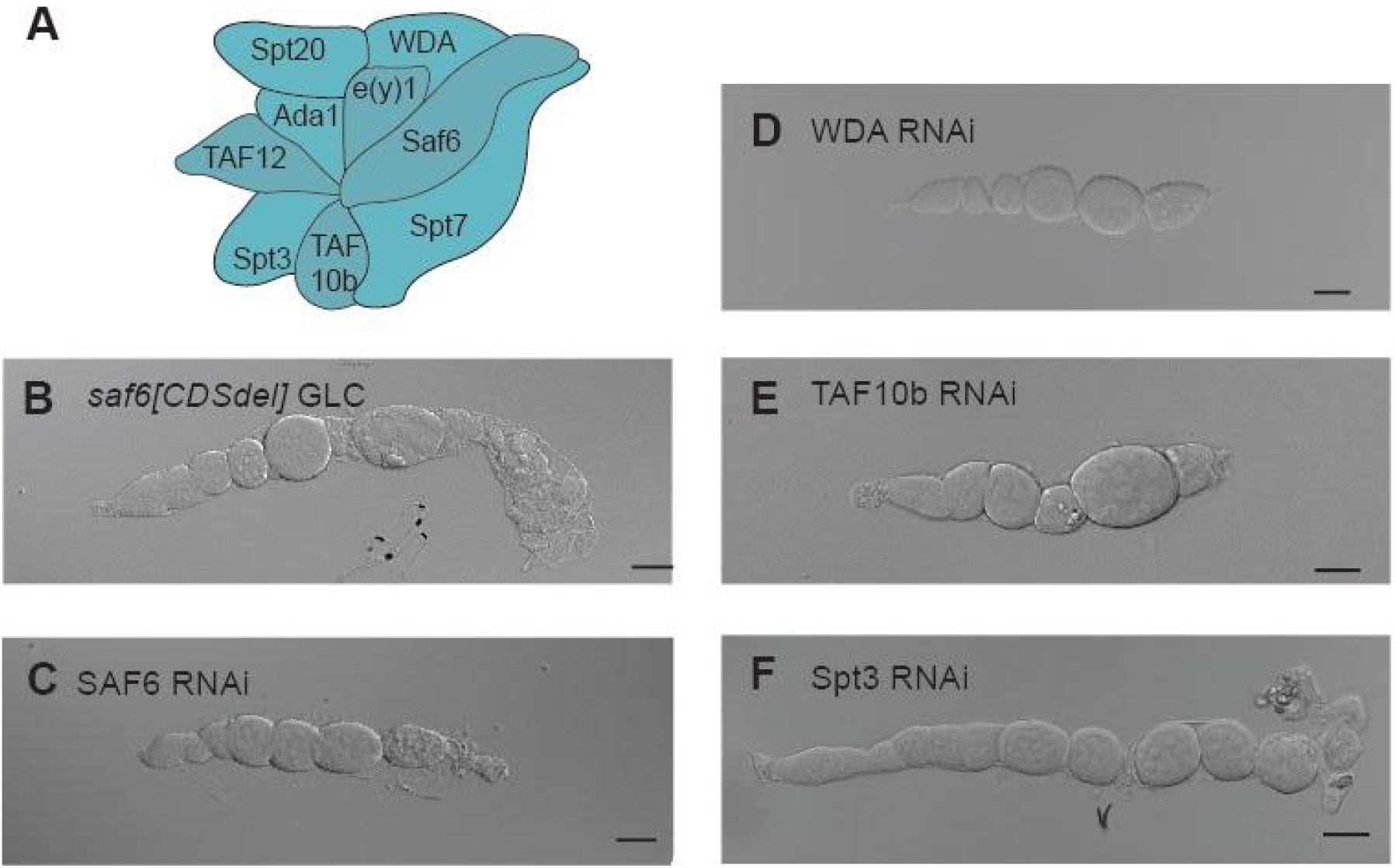
The core module of the SAGA complex is required for mid-oogenesis. (A) Overview of SAGA core subunits. (B) *saf6[CDSdel]* ovariole. (C) SAF6 RNAi ovariole. (D) TAF10b RNAi ovariole. (E) Spt3 RNAi ovariole. Scale bar: 50 µM

### SAGA core subunits may function through TBP

Based on the homology with yeast SAGA complexes [22, 45], we infer that WDA interacts with SAF6 and TAF10b, which are important for the structural configuration of Spt3 that allows the binding of TBP. Therefore, we hypothesized that the SAGA-specific TAF subunits are critical for oogenesis because they interact with TBP at the promoter.

To test this, we targeted Spt3 by RNAi in the germline. This also arrested oogenesis at stage 6-7 and produced a phenotype highly reminiscent of those observed for the TAFs (Fig 2F). The number of immature egg chambers increased, but they remained round, and the oocyte did not enlarge at the distal end. This result suggests that the SAGA-specific TAF subunits are required for the function of Spt3 and thus likely influence the deposition of TBP onto promoters during oogenesis.

We next tested whether the direct depletion of TBP in the germ line caused a similar phenotype. However, germ line depletion of TBP led to a more severe phenotype (Fig 3A), indicating that other TBP-containing complexes function in the germ line. To test this, we depleted TAF1 (Fig3 B) and TAF4 (Fig 3C), which are TFIID-specific TAF subunits. Depletion of TAF4 and TAF1 all caused severe agametic phenotypes, confirming that TBP and TFIID play a broader role during oogenesis than the SAGA complex, and agrees with other studies in yeast demonstrating that a smaller gene set is affected by loss off SAGA versus TFIID. [19].

**Fig 3.**
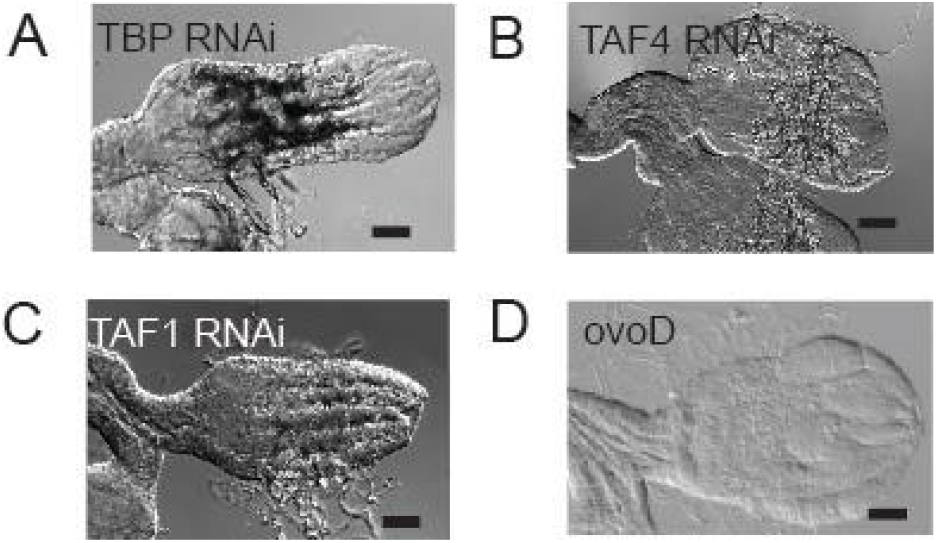
TFIID subunit depletion blocks oogenesis earlier than SAGA subunit depletion. (A) TBP RNAi ovary pair (B) TAF4 RNAi ovary pair. (D) TAF1 RNAi ovary pair. (E) OvoD control ovary pair. (F) Scale bar: 50 uM.

### The SAGA complex broadly binds to various promoter types

The stronger phenotype associated with depletion of TBP compared to SAGA-specific core subunits prompted us to test whether the role of SAGA is specific for certain promoter types. Such distinction is of particular interest in a developmental contexts, where promoter types may be preferentially used [46-48]. For example, TATA promoters in *Drosophila* are enriched amongst the earliest expressed developmental genes in the embryo [20, 49], and previous studies suggested that TATA promoters have differential requirements for TFIID and SAGA {Huisinga, 2004 #99, 50]. Indeed, the phenotype of Spt3 depletion pointed to a connection between SAGA and TBP, which engages the classic TATA promoters with high affinity [51].

To determine how SAGA occupies promoters in ovary tissue genome-wide, we performed chromatin immunoprecipitation followed by sequencing (ChIP-seq) using antibodies against the SAGA core module subunits WDA, SAF6 and Spt3, and against Ada2b, a SAGA HAT module subunit. We then compared their occupancy patterns and analyzed how they relate to the promoter activities. As measurement for each promoter’s activity, we used Cap Analysis Gene Expression (CAGE) data from ovaries of virgin females (modENCODE_5368 [52], which yielded capped transcripts for approximately 6,000 promoters (> 2 TPM).

Our data revealed widespread occupancy of all SAGA subunits at active promoters, as well as a strong correlation between the different subunits (Fig 4). When we sorted the top 4,000 active promoters by WDA occupancy, the occupancy pattern was visually very similar for the other subunits (Fig 4A). For all subunits, the average binding pattern was strongest near the TSS, supporting its function at the core promoter (Fig 4B). To investigate the relationship between occupancy and transcription, we grouped all active genes into ten quantiles of CAGE expression levels and calculated within each quantile the binding levels of SAGA at each promoter. With decreasing expression levels, we observed a gradual decrease in occupancy for all examined SAGA subunits (Fig 4C). These results show that SAGA binding in the ovaries is widespread at active promoters and that the degree of occupancy correlates with transcriptional activity.

**Fig 4.**
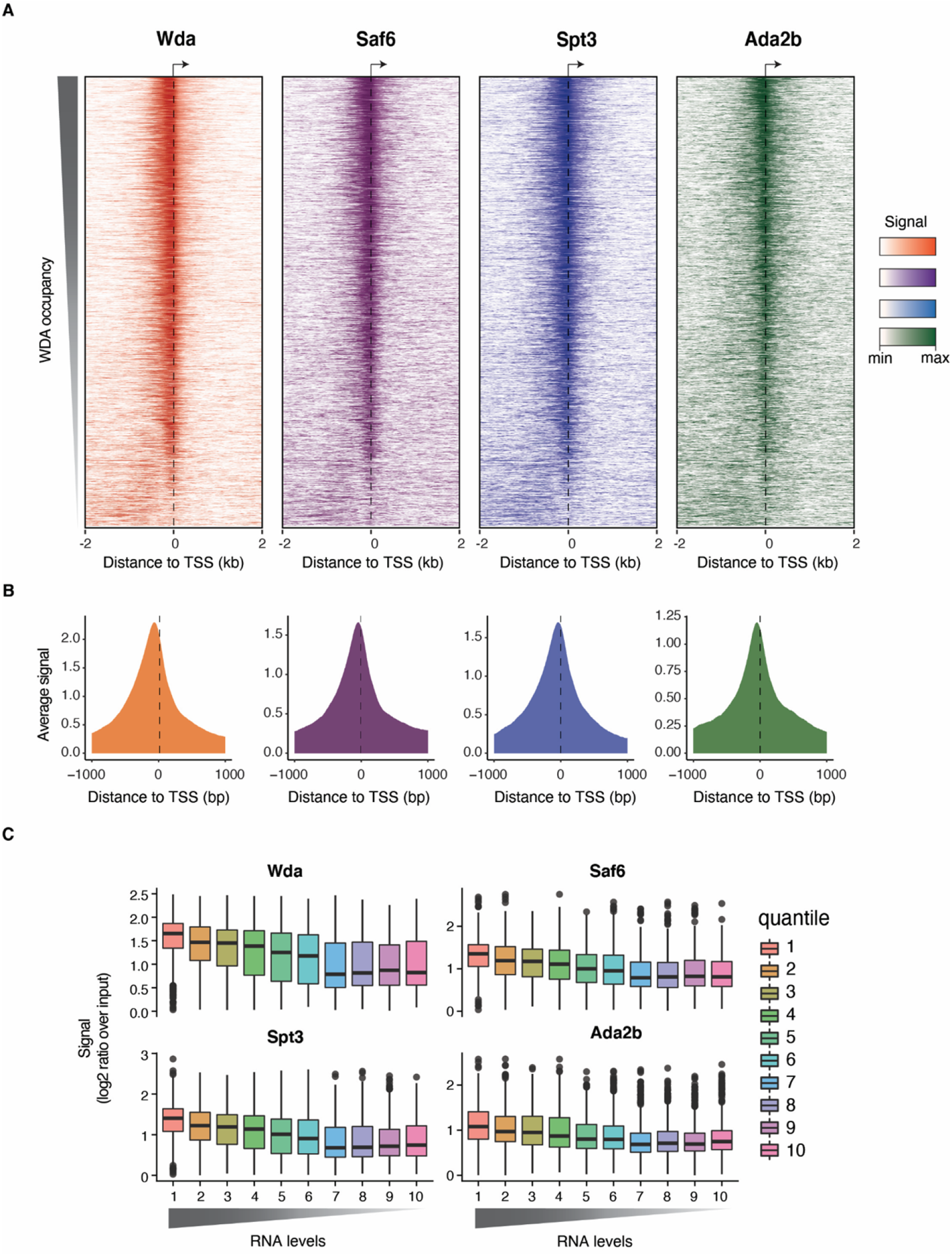
Genome-wide occupancy of SAGA subunits in ovary tissue shows that SAGA broadly binds to active promoters. (A) Normalized ChIP signal in a 1001 bp window centered at the TSS of the top 4,000 genes with the highest expression in ovary tissue. Promoters were sorted by WDA signal in all 4 panels. (B) Metagene plots showing the average binding at these 4,000 promoters for each SAGA subunit. (C) The 4,000 active genes were divided into ten quantiles based on CAGE expression levels (decreasing from left to right) and each box plot panel displays the SAGA subunit occupancy per quantile (log2 ratio over input). This shows that SAGA occupancy levels correlate with transcriptional activity.

We next analyzed whether the ChIP-seq occupancy of the SAGA subunits differs between promoter types. We analyzed the following *Drosophila* promoter types: 1) TATA promoters, 2) promoters with Downstream Promoter Region (DPR) elements, 3) promoters with the polypyrimidine initiator element TCT, and 4) housekeeping (HK) promoters that may contain a DRE, Ohler motifs 1, 6 and 7 (but lack TATA, TCT or DPR elements) [53-55]. TCT promoters require TBP Related Factor 2 (TRF2) and not TBP for their activation [56]. TRF2 is structurally similar to TBP, but lacks high affinity binding for TATA boxes [28, 57, 58] and binds to different sites in the *Drosophila* germ line [30]. In addition, TRF2 interacts with DREF, the factor that binds to the DRE motif [28, 30, 59]

We identified 124 TATA genes, 142 TCT genes, 60 DPR genes and 856 HK genes among the active promoters with CAGE-seq (S2). Using this promoter classification, we found that SAGA occupancy was not higher at TATA promoters or lower at promoters that preferentially use TRF2, arguing against promoter-specific recruitment of SAGA.

ChIP-seq data are however of low resolution and may capture unspecific binding. In contrast, our established ChIP-exo protocol called ChIP-nexus has higher specificity and near base-resolution [43]. It has previously revealed the exact locations of pre-initiation complex components along the promoter, including TBP [60]. Since we could not perform ChIP-nexus experiments in ovaries, we used *Drosophila* Kc167 cells and performed ChIP-nexus experiments against the four SAGA subunits, as well as against TBP and TRF2 (Fig 5 and S4). To analyze their binding profile for each promoter type, we identified 164 TATA genes, 103 TCT genes, 320 DPR genes and 1385 HK genes among active genes (S2).

**Fig 5.**
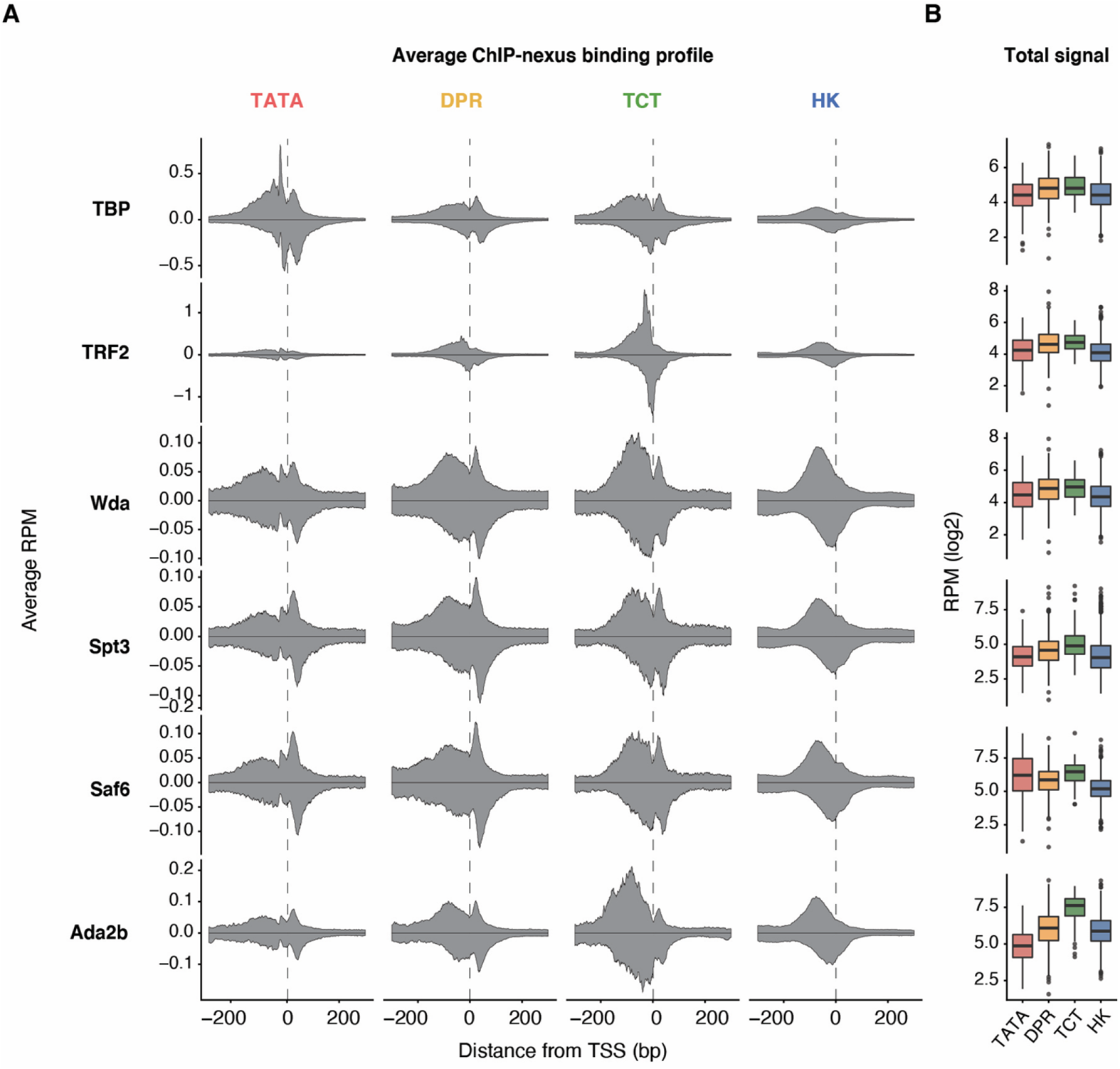
SAGA footprint at different promoter types in cells. (A) Average footprints of SAGA subunits in reads per million (RPM) for clusters of genes with different promoter types (left to right TATA, DPE-like, TCT and HK genes). From top to bottom SAGA subunits Spt3, SAF6, WDA and Ada2b, TBP and TRF2. Grey horizontal dashed lines mark 0.1 average RPM. Blue horizontal dashed lines mark 1 average RMP. (B) Boxplot displaying total signal per promoter type.

**Fig 6.**
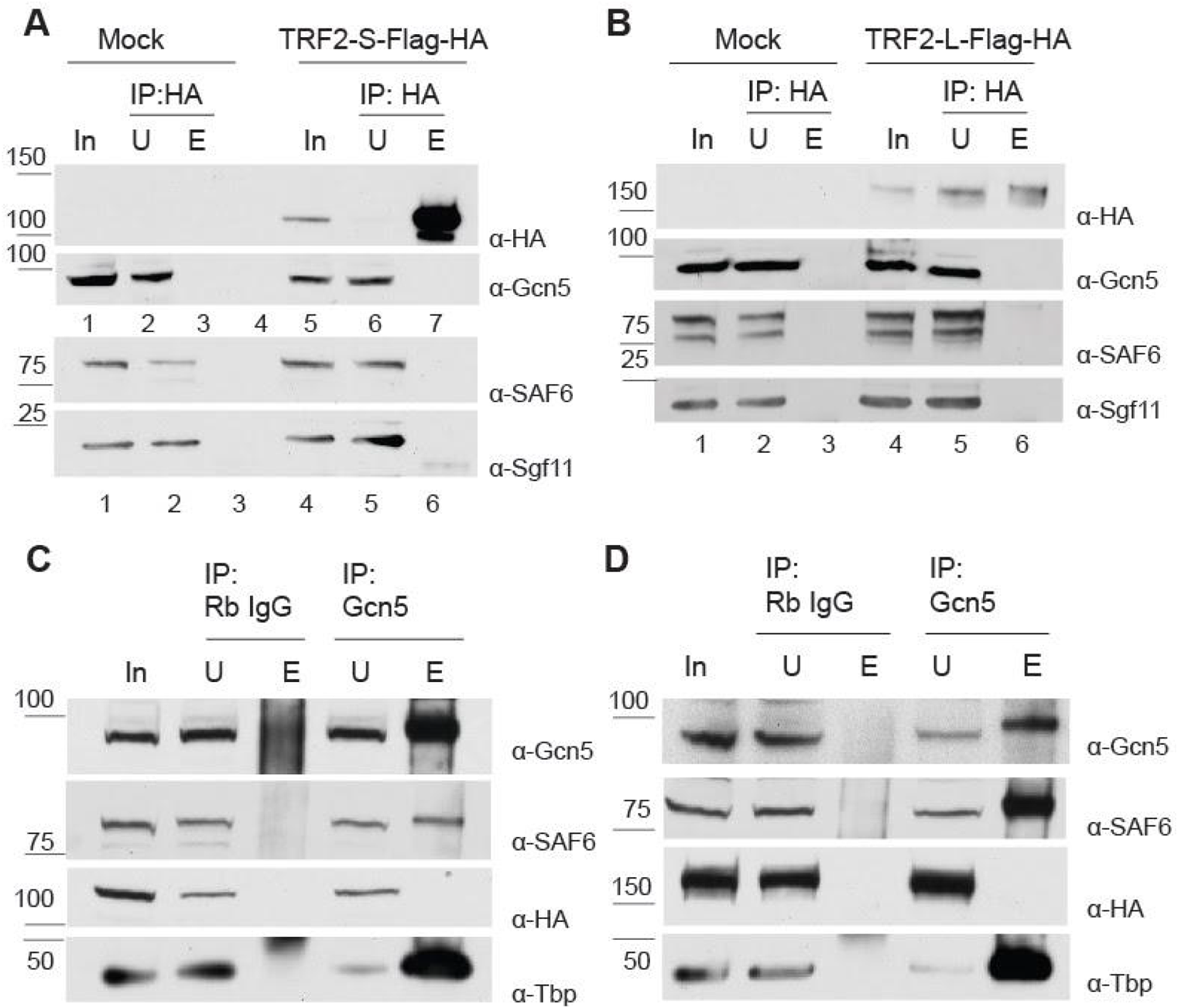
SAGA and affinity-tagged TRF2 isoforms do not stably interact in S2 cells. (A) Reciprocal coimmunoprecipitation after transient transfection with pAcTRF2-S-Flag-HA. Using rat anti-HA to immunoprecipitated TRF2-S-Flag-HA, SAGA did not coimmunoprecipitate. (B) Reciprocal coimmunoprecipitation after transient transfection with pAcTRF2-L-Flag-HA. Using rat anti-HA to immunoprecipitated TRF2-L-Flag-HA, SAGA did not coimmunoprecipitate. (C) Coimmunoprecipitation after transient transfection with pAcTRF2-S-Flag-HA. Using Gcn5 as a bait, affinity-tagged TRF2-S does not coimmunoprecipitate. (D) Coimmunoprecipitation after transient transfection with pAcTRF2-L-Flag-HA. Using Gcn5 as a bait, affinity-tagged TRF2-L does not coimmunoprecipitate.

As expected, the highest levels of TBP were observed at TATA promoters (Fig 5). TRF2 had not previously been analyzed with ChIP-nexus, but the high levels of TRF2 at the TCT and HK promoters were consistent with previous ChIP experiments and functional studies {Hochheimer, 2002 #1462; Wang, 2014 #1480. In contrast to TBP and TRF2, the four SAGA subunits appeared to be enriched at comparable levels across all four promoter types similar to the ovary tissue (S3, S4).

At this level of resolution, it became apparent that there are slight differences in the SAGA binding patterns between promoter types. Each SAGA subunit binds certain locations upstream and downstream of the TSS with variable amounts. For example, SAGA subunits bound mainly upstream of the TSS at TCT and housekeeping genes, but appeared more prominent downstream of the TSS at TATA and DPR genes (Fig. 5). Since SAGA is recruited by activators (reviewed in {Soffers, 2020 #1760;Chen, 2021 #1837}), this difference could be due to the different location of the enhancer regions. These are located directly upstream of the TCT and housekeeping promoters, but are found more distally at TATA and DPR genes and thus would not contribute to the signal in the average profile around the promoter. While these profiles with upstream and downstream binding were interesting, they did not support our hypothesis that SAGA is specifically associated with TBP at TATA promoters.

### SAGA binds TBP robustly but was not found to bind TRF2

Since SAGA was strongly bound at TCT genes, where TRF2 functions, we considered that SAGA may also be associated with TRF2. We therefore tested if the SAGA complex stably interacts with TRF2 by co-immunoprecipitation. We transiently transfected epitope-tagged constructs in S2 cells. We tested both TRF2 isoforms since both form higher molecular weight complexes [30, 59]. However, we did not detect SAGA subunits after immunoprecipitation of TRF2 (Fig 7 A, B). Likewise, we failed to detect TRF2 isoforms after immunoprecipitation of the SAGA complex (Fig 7 C, D). As control, endogenous TBP interacted robustly with SAGA subunits (Fig 7 C, D). These results indicate that epitope-tagged TRF2 and SAGA do not stably interact, consistent with similar observations in murine testes, where TRF2 co-immunoprecipitated with TFIIA but not TAF7L, although it co-localized in ChIP experiments with both [31]. It is therefore possible that the colocalization of SAGA and TRF2 *in vivo* requires a chromatin template.

**Fig 7:**
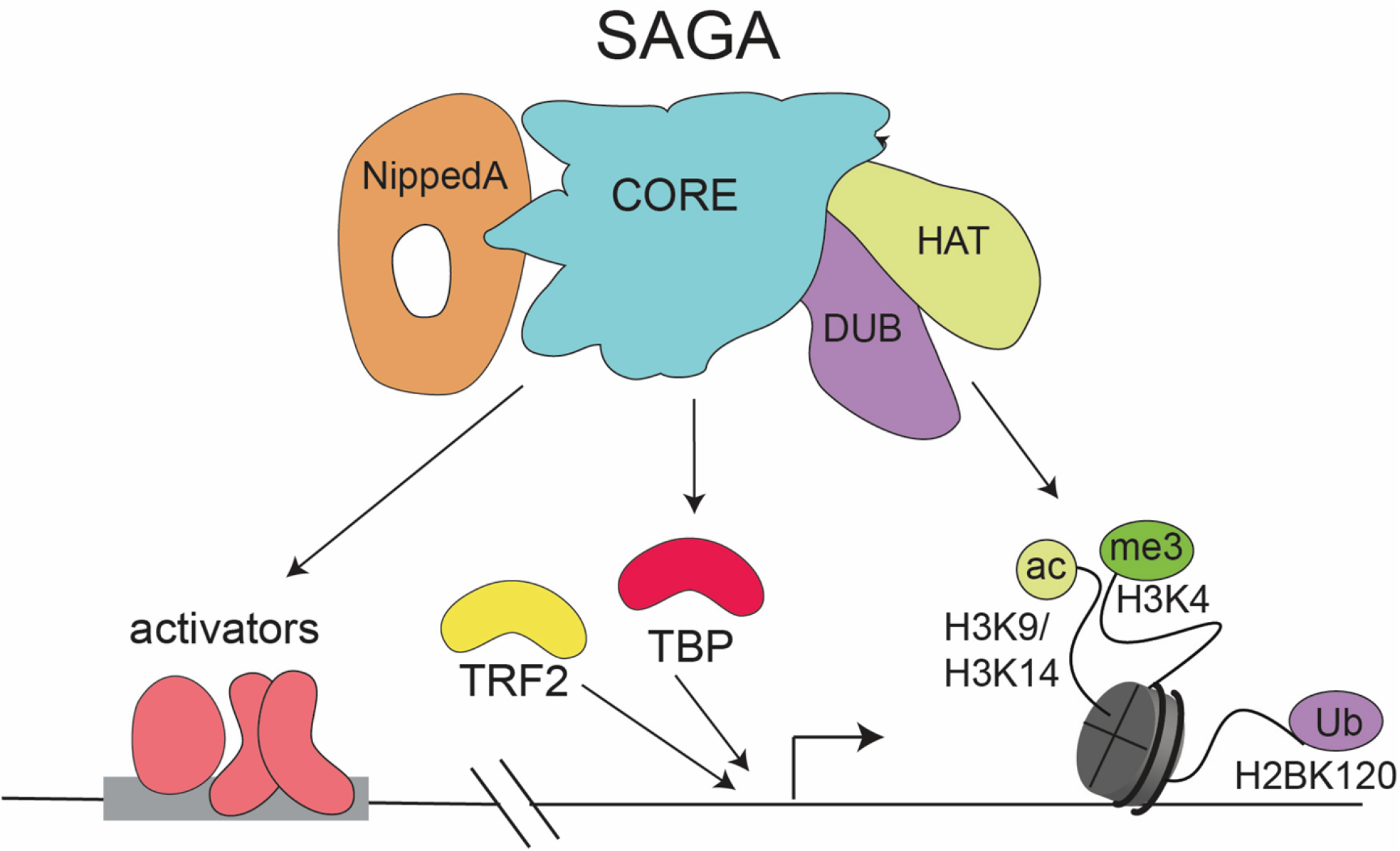
Cartoon of SAGA core module functions during oogenesis. The SAGA complex binds upstream and downstream of the TSS at active genes, without discriminating by promoter type. The upstream peak is likely due to interactions of the core module and Nipped A, which interact with activators. The complex is also present at the +1 nucleosome, where it can read H2bUb, H3K4me3 and H3K9/4ac and is retained. After its recruitment to the +1 nucleosome, the enzymatic DUB and HAT activities of SAGA retain the complex, carryout histone modifications and may help maintain the core promoter region in an open state, while the core module may contribute to deposition of TBP onto the core promoter. We speculate that through its capacity to interact with TBP, the SAGA core module may help to increase the concentration of TBP near promoter chromatin.

## Discussion

SAGA is a large versatile coactivator complex in which all subunits and modules typically tightly interact with each other biochemically and where the modules function synergistically to activate transcription. We have hence considered SAGA biochemically to be one entity and expected that the disruption of any part would disrupt the functionality of the entire complex. However, we found that the genetic phenotype differs between SAGA modules during *Drosophila* oogenesis. Our present work demonstrates that the depletion of the SAGA core module arrests development during mid-oogenesis, while our previous work showed that depletion of the HAT module produced a milder phenotype and the DUB module was not required at all for oogenesis [25]. Together, these data suggest that not all SAGA modules are equally important in this developmental context.

Such differential genetic requirements of subunits or modules have sometimes been interpreted as the result of their differential recruitment to specific genes [25, 26]. However, our ChIP data suggest that all subunits and modules are recruited to all active promoters, consistent with the strong biochemical integrity of the SAGA complex under normal conditions. Moreover, we found that the occupancy of SAGA was not promoter-specific. This is surprising since our genetic and biochemical data suggest a specific interaction of SAGA with TBP, but not TRF2. These results strengthen the more recently established role of SAGA as general transcription coactivator [1, Hahn 2020, 61] and support the idea that the different functions of SAGA are not reflected in differential recruitment to chromatin.

Since the genetic analysis nevertheless pointed to functional differences among the modules, we propose the handyman principle for the function of large biochemical complexes such as SAGA. While the correct functioning of a house requires many components, certain components break more easily and require more attention by the handyman, giving the impression that they are more essential. Likewise, gene regulation occurs in a redundant, buffered fashion by multiple complexes and modules, but under specific circumstances, a single regulatory module or enzymatic activity may become rate-limiting. Such a model is consistent with previous observations where depletion of an essential enzyme or transcription factor such as P-TEFb in early *Drosophila* embryos only affected a few highly expressed genes in an apparent unspecific fashion [62].

Consistent with the handyman principle, our data during oogenesis suggest that the core module’s promoter function is most critical, although all modules are recruited to all active promoters. This could be because of its interaction with TBP, which helps position TBP upstream of the transcription initiation site, although SAGA may also function at promoters that primarily use TRF2 for activation (Fig 7).

Another explanation for the critical requirement of the core module is that it mediates the recruitment of other SAGA modules to chromatin. The recruitment and activity of SAGA depends on activator proteins and the core module bridges Nipped-A (yeast Tra1), the major hub for activator interactions to the enzymatic modules. Moreover, the module itself also contains subunits directly involved in promoter binding [63]. Consistent with this idea, histone acetylation by the SAGA HAT module depends on promoter recruitment [2, 8] and this requires certain core module subunits in *Drosophila*[42].

Our high-resolution ChIP-nexus data provided additional insights into the interaction of SAGA subunits with chromatin. While previous ChIP data only showed broad associations with promoters and RNA polymerase II [25, 64]. ChIP-nexus detected preferential binding in regions upstream of the start site, where activators are found [55], consistent with the role of activators in SAGA recruitment. In addition, the ChIP-nexus data revealed binding of SAGA downstream of the start site, near the +1 nucleosome.

This raises the interesting possibility that SAGA binding may be stabilized through its interaction with nucleosomes. Several biochemical studies support this possibility. The bromodomains within the catalytic subunits of SAGA interact with acetylated histones [65]and the tandem tudor-domains within the HAT module serve as H3K4me3 readers, whose levels depend on H2Bub, to further stimulate the processive acetylation by SAGA [66]. Thus, there appears to be extensive crosstalk, in which SAGA’s ability to recognize its own histone modifications may promote further interactions with nucleosomes. Given this cross-talk, we propose a model in which SAGA’s occupancy at promoters is in part due to the recruitment via activators, and in part due to its retention on chromatin via modified nucleosomes.

## Materials and methods

### Antibody production

Affinity purified antibodies for ChIP analysis were generated by the GenScript Company following the PolyExpress Premium Antigen-Specific Affinity Purification Production protocol. Antigens included full length Gcn5, WDA and SAF6.

### Fly housing and ovary dissection

*Drosophila* melanogaster were raised and maintained at 25°C with a 12 hour/12 hour, light/dark cycle in non-crowding conditions. The animals were reared at 25°C on standard fly food containing 14 g inactive dry yeast (Genesee Scientific, Cat. Num. 62-108), 46 g yellow corn meal (VWR, Cat. Num. 75860-346), 6.9 g agar (Genessee Scientific, Cat. Num. 6-103), 11.5 mL corn syrup (Reinhart Food Service, Cat. Num. 31494), 53.5 g dried malt (Briess Industries Inc, Cat. Num. 5728), 8mL 30% Tegosept (Genesee Scientific, Cat. Num. 20-258) in ethyl alcohol (Sigma-Aldrich, Cat. Num. 459836), 6 mL propionic acid (Sigma-Aldrich, Cat. Num. 402907), and 900 mL of water. Ovaries were dissected in M3 Shields and Sang media from females conditioned three days on apple juice plates supplemented with yeast paste. A list of genotypes is included in S1 Table.

### Cell culture and transient transfections

Wild type S2 or Kc167 S2 cell lines were maintained at 1-2 e^6^ cells/mL in SFX medium (HyQ SFX-Insect;HyClone Laboratories, Inc) at 27°C. Transient Transfections of S2 cells were performed using Qiagen Effectene Transfection reagent (Qiagen 30425) at a 1:25 DNA-Enhancer ratio. With respect to cell density and culture volume, 3e^6^ S2 cells resuspended in 2.5 mL SFX medium were transfected with 0.4 µg plasmid DNA and this was scaled up when required. pAcTRF2-S-Flag-Ha and pAcTRF2-L-Flag-Ha were kindly gifted by the Gershon laboratory [53].

### Coimmunoprecipitation

Nuclear extracts of 15 e^7^ S2 cells were prepared five days post-transfection. Cells were collected by 1 800 x g centrifugation for five minutes at 4°C. Cells were next resuspended in 1mL lysis buffer (10 mM HEPES [pH 7.9], 1.5 mM MgCl_2_, 10 mM KCl and 1% NP-40) supplemented with 1mM PMSF (Millipore Sigma) and 1x Roche cOmplete Protease Inhibitor Cocktail Tablets (Millipore Sigma). After five minutes lysis on ice the nuclei were collected by centrifugation at 10 400 x g for five minutes at 4°C. The nuclei pellet was incubated with 0.5 mL high-salt Nuclear Extraction buffer (20 mM HEPES [pH 7.9], 25% glycerol, 1.5 mM MgCl_2_, 0.2 mM EDTA, 0.1% Triton-X100 and 420 mM NaCl) supplemented with 5 µl benzonase (Millipore Sigma), 1mM PMSF and 1X protease inhibitor for one hour at 4°C while rotating. The soluble nuclear proteins were collected as supernatant after centrifugation at 15 000 RPM for 20 minutes at 4°C. Protein concentrations were measured by Bradford assay.

Immunoprecipitations were performed with 1.5 mg nuclear extract diluted to 0.15 M NaCl with Dignam buffer. Each IP reaction was performed with 50 µl Dynabeads precoated with 1 µg of protein. The capturing antibody (rabbit FL Gcn5 (Genscript), rat high affinity anti HA (MilliporeSigma 11867423001)) or IgG (Santa Cruz) were incubated with 50 µl washed Dynabeads resuspended in 1 mL PBS (A for rabbit Invitrogen 10001D/, G for rat antibodies Invitrogen10003D). After binding for two hours at 4°C while rotating, unbound bait protein was removed, and the beads were washed one time in 1mL cold PBS prior to incubation with the nuclear lysate for immunoprecipitation. The next morning, the unbound extract was collected, and the beads were washed three times in 1 mL wash buffer (10 mM HEPES [pH 7.9], 1.5 mM MgCl_2_, 150mM NaCl, 10 mM KCl, 0.2% Triton X-100) prior to elution in 100µl 2x Laemmli Sample buffer (Bio-Rad160747). Western blots were performed with 20 µl input (equaling 20 µg protein), 20 µl unbound and 10-30 µl eluate.

### CRISPR/CAS9 Generation of SAF6 mutant fly lines

A SAF6 CDS deletion line was generated by targeting the 5’ and 3’ UTR by a guide RNA and recombination of a DsRed cassette with the CDS. The 5’ gRNA (5’ GATTATTAACCCAACTATCGA 3’) (primers 1 and 2, S2 Table) and 3’ guide RNA (5’ GAGTAAAGTTGTATCACCTTT 3’) (primers 3 and 4, S2 Table) were each cloned into the BbsI restriction site of pBFv-U6.2 as described before [67]. Amplicons of 500 bp spanning the guide region were generated using primers 5-8 to test if the reported Flybase genome sequence was accurate and the parent fly (BL55281) was free of SNPs.

A donor construct was created that contained a DsRed cassette flanked by 1KB homology arms that matched the upstream and downstream 1Kb DNA sequence of each guide (primers 9-12, S2 Table). The DsRed cassette and pHSG289 backbone was PCR-amplified from the pHDScarless DsRed plasmid (https://flycrispr.org/scarless-gene-editing) and the homology arms were PCR-amplified from wild-type fly genomic DNA. The PCR products were combined via HiFi DNA assembly (NEB). The plasmid was validated using Sanger sequencing.

A total of 300 embryos of parent fly (BL55821) were injected at SIMR by Paul Leal. All surviving adults were mated to w[1118]/Dp(1;Y)y[+]; noc[Sco]/SM6a. The F1 larvae were screened for DsRed signal and all adult males with DsRed positive eyes mated in single-male crosses w[1118]/Dp(1;Y)y[+]; noc[Sco]/SM6a. Male F2 progeny with DsRed balanced by Sm6A were again mated to this balancer create stable stocks of the genotype w[118];saf6CDSdelDsRed/SM6a, abbreviated as saf6[CDSdel].

### WDA allele EMS mutagenesis

The wda[EMS] stock was generated by a ethyl methanesulfonate (EMS) screen performed by Xuanying Li and Susan Abmayr. Adult males (P{neoFRT}82B ry[506]) were starved at 3-5 days post-eclosion for three hours and transferred to vials saturated with 30mM EMS, 100mM Tris-HCl and 10% sucrose. After 20 hours, the males were transferred to fresh food for three changes to clear the EMS and mated to virgin females (y,w;D[3],gl[3]/TM3,Tb[1],Ser[1]) 2-6 days post-eclosion. Progeny TM3 males were mated to WDA[4] and WDA[8] [42] and non-complementing alleles were recovered. (genotype: p{neoFRT}82B ry[5-6,wda*/ TM3,Tb, Sb]). Non-complementation crosses were performed again using deficiency stock BL25694(w[1118]; Df(3R)BSC619/TM6C, cu[1] Sb[1]. A total of 46 stocks did not complement the deficiency. The balancer was exchanged to TM3,p{w[+mC]=GAL4-tw.G}2.3,p{UAS-2EGFP}AH2.3,Sb[1],Ser[1]. Genomic DNA was obtained from homozygous embryos. Mutation in WDA was confirmed by PCR (primer 13 and 14, S2 Table) and sequencing. This allele contains a 321 bp deletion in exon1 through exon 2. Random mutations were removed by outcrossing the mutagenized chromosome for two generations to Oregon R before the phenotypic analysis.

### Germ line clones

To create germline clones, one needs a recombinant stock that contains an FRT site that is located at the same arm as where the mutated allele is positioned. (FRT82B for WDA or chromosome 3R, and FRT40A for SAF6 on chromosome 2L). The saf6 mutant allele was created in BL55821 and FRT^40A^ was recombined into stock w[118];saf6[CDSdel]/SM6a. At the same time, a heat-shock flip was introduced on the X-chromosome using y,w,HSFLP;FRT^40A^-ubiGFP/CyO since the corresponding ovo^D^ stock does not carry one (Bl2121;P{w[+mC]=ovoD1-18}2LaP{w[+mC]=ovoD1-18}2Lb {ry[+t7.2]= neoFRT}40A/ Dp(?;2) bw[D], S[1] wg[Sp-1] Ms(2)M[1] bw[D]/CyO) and balanced with y,w,HSFLP;Sco/CyO.

The actual germ line clones were created by crossing virgins from each “mutation, FRT stocks” to the corresponding ovoD males to introduce recombination. Larvae were heat-shocked for 2hr at 37°C for two days starting from the second day after hatching. Homozygous virgins were collected and used for further analyses.

### Germ-cell specific knockdown in ovaries

For each protein of interest, we tested where possible the efficacy of multiple RNAi sequences expressed from either the second or third chromosomes using pValium20 and pValium22 lines where possible. The knockdown was tested at 25 and 29 °C. UAS RNAi lines to target WDA, SAF6 or TAF10b were generated by Rainbow Genetics Inc (S1 Table). To induce expression of the RNAi, UAS-RNAi flies were crossed to the maternal triple driver was chosen that expressed GAL4 maternally, with three promoters, and targets the germ line. This is stock BL31777, genotype P{w[+mC]=otu-GAL4::VP16.R}1, w[*]; P{w[+mC]=GAL4-nos.NGT}40; P{w[+mC]=GAL4::VP16-nos.UTR}CG6325[MVD1] and throughout the manuscript referred to as MTD-Gal4. Throughout the text the SAGA subunit RNAi/MTD-Gal4 genotype is abbreviated: e.g. UAS RNAi e(y)1 (BL 32345)/MTD-Gal4 is refer to as E(y)1 RNAi.

### Regular ChIP in ovaries

ChIP was performed on ovaries as described before [68]. To minimize variation in the ovary ChIPs three batches of chromatin were prepared and all subunit IPs were performed on the same batch of chromatin using 20 ug chromatin to generate three biological replicates. ChIP for WDA and SAF6 was performed with 10 µg antibody, Ada2b, Spt3 and Sgf11 with the amounts described in [25]. Data was aligned to *Drosophila* genome version dm6, using bowtie with parameters -k 1 -m 3. Gene definitions utilize Ensembl 87. Peaks were called using MACS2 under default parameters. Peaks present in at least two of three replicates were combined and reduced to create a reference peak set for potential subunit binding locations.

### ChIP-nexus in cells

For each ChIP-nexus experiment, 10^7^ Kc167 cells were fixed with 1% formaldehyde in SFX culture media at room temperature for 10 minutes with rotation. Fixed cells were washed with cold PBS, incubated with Orlando and Paro’s Buffer (0.25% Triton X-100, 10 mM EDTA, 0.5 mM EGTA, 10 mM Tris-HCl pH 8.0, supplemented with 1x Protease Inhibitor) for 10 minutes at room temperature with rotation, and then centrifuged and re-suspended in ChIP Buffer (10 mM Tris-HCl, pH 8.0; 140 mM NaCl; 0.1% SDS; 0.1% sodium deoxycholate; 0.5% sarkosyl; 1% Triton X-100, supplemented with 1x Protease Inhibitor). Sonication was performed with a Bioruptor Pico for five rounds of 30 seconds on and 30 seconds off. Chromatin extracts were then centrifuged at 16000 g for 5 minutes at 4°C, and supernatants were used for ChIP.

To couple Dynabeads with antibodies, 50 µl Protein A and 50 µl Protein G Dynabeads were used for each ChIP-nexus experiment and washed twice with ChIP Buffer. After removing all the liquid, Dynabeads were resuspended in 400 µl ChIP Buffer. 10 µg antibodies were added, and tubes were incubated at 4°C for 2 hours with rotation. After the incubation, antibody-bound beads were washed twice with ChIP Buffer.

For chromatin immunoprecipitation, chromatin extracts were added to the antibody-bound beads and incubated at 4°C overnight with rotation and then washed with Nexus washing buffer A to D (wash buffer A: 10 mM Tris-EDTA, 0.1% Triton X-100, wash buffer B: 150 mM NaCl, 20 mM Tris-HCl, pH 8.0, 5 mM EDTA, 5.2% sucrose, 1.0% Triton X-100, 0.2% SDS, wash buffer C: 250 mM NaCl, 5 mM Tris-HCl, pH 8.0, 25 mM HEPES, 0.5% Triton X-100, 0.05% sodium deoxycholate, 0.5 mM EDTA, wash buffer D: 250 mM LiCl, 0.5% IGEPAL CA-630, 10 mM Tris-HCl, pH 8.0, 0.5% sodium deoxycholate, 10 mM EDTA). End repair and dA-tailing were performed using the NEBNext End Repair Module and the NEBNext dA-Tailing Module. ChIP-nexus adaptors with mixed fixed barcodes (CTGA, TGAC, GACT, ACTG) were ligated with Quick T4 DNA ligase and converted to blunt ends with Klenow fragment and T4 DNA polymerase. The samples were treated with lambda exonuclease and RecJ_f_ exonuclease for generating Pol II footprints at high resolution. After each enzymatic reaction, the chromatin was washed with the Nexus washing buffers A to D and Tris buffer (10 mM Tris, pH 7.5, 8.0, or 9.5, depending on the next enzymatic step).

After RecJ_f_ exonuclease digestion, the chromatin was eluted and subjected to reverse crosslinking and ethanol precipitation. Purified single-stranded DNA was then circularized with CircLigase, annealed with oligonucleotides complementary to the BamHI restriction site and linearized by BamHI digestion. The linearized single-stranded DNA was purified by ethanol precipitation and subjected to PCR amplification with NEBNext High-Fidelity 2X PCR Master Mix and ChIP-nexus primers. The ChIP-nexus libraries were then gel-purified before sequencing with Illumina NextSeq 500.

### Promoter type analysis

TSSs from non-overlapping Flybase protein coding genes (fb-r5.47) were re-annotated using CAGE-seq data from *Drosophila* melanogaster kc167 cells (<u>modENCODE_5333</u>) and virgin fly ovaries (modENCODE_5368). For TSS tag clustering (TCs), replicates were merged (only for Kc 167cells), and low-quality reads were removed using the CAGEr package from R. Clusters with only one TSS (singletons) were discarded unless the normalized signal (TPM) was equal or above 2. TSSs within 40-bp of each other were clustered together and only clusters with at least 1 TPM from all TSS positions were considered expressed and kept for further analysis. To separate focused from dispersed promoters, the width of TCs was calculated as described by [69]. Promoters with a TC width <= 9-bp were considered to have a focused transcription initiation pattern while promoters with a TC width > 9-bp were included in dispersed category.

Promoter types were further defined by the presence or absence of the motifs listed in S3 Table. TATA and TCT promoters were only required to contain the TATA-box and the TCT element, respectively, but were not excluded for having other elements. Conversely, DPR and housekeeping promoters were additionally filtered to not have motif elements from any other of the promoter categories. Thus, DPR promoters were allowed the presence of MTE, DPE and PB but not TATA, TCT, Ohler1/6/7 or DRE, whereas housekeeping promoters were allowed to contain Ohler1/6/7 or DRE but not TATA, TCT, MTE, DPE or PB.

DNA sequences at each promoter category (TATA, TCT, DPR and Housekeeping) were obtained from the dm6 genome and represented as heatmap. The motifs logos were generated using the R package ggseqlogo. ChIP-nexus average binding profiles (metaprofiles) were plotted in 201-bp window using the average footprints for each factor in reads per million (RPM) on the positive strand (above line) and negative strand (below line) across the different promoter groups. Total signal distributions (boxplots) were calculated at each promoter as the total number of reads (RPM) in a 101-bp window centered at the TSS for ChIP nexus data and 501 for regular ChIP-seq data.

### Differential Contrast Imaging

Most DIC images were obtained using a Nikon Ti2 motorized widefield microscope equipped with Nikon Perfect Focus technology and a sCMOS camera. However, due to COVID-19 research restrictions selected images had to be captured using an Axioplan2 wide field microscope using a 20x air objective. Multiple images were captured to cover each ovariole and merged with Photoshop using the default settings (Files□ Automate □photo merge). Grey boxes have been placed behind the image to obtain equal panel sizes when required.

### Western blotting analysis

The gel filtration samples were resolved on 10% SDS-PAGE gels. The HAT assay samples were resolved on 15% SDS-PAGE gels. Gels were transferred to a PVDF membrane and blocked for one hour at 4°C in 5% milk in Tris-buffered saline (TBS) and 0.1 % Tween-20. Primary antibodies were diluted in 5% milk in TBS and 0.1%Tween-20 and incubated overnight at 4°C. The following antibodies were used: Gcn5 (rabbit polyclonal, 1:1000, (GenScript antibody services, Atlanta, GA, USA anti full-length Gcn5); Ada2b (rabbit polyclonal, 1:1000; GenScript anti-amino-acid 1-330); anti-Flag-horseradish peroxidase (mouse, 1:5000; Sigma Millipore); H3 (rabbit, 1:10000; Abcam); H3K9/K14ac (rabbit, 1:1000, Sigma Millipore); Donkey anti-rabbit IgG-horseradish peroxidase (1:5000, Fisher Scientific). The rabbit TBP antibody was kindly shared by the Zeitlinger laboratory.

The data discussed in this publication have been deposited in NCBI’s Gene Expression Omnibus [70] and are accessible through GEO Series accession number GSE166224.

## Acknowledgements

We thank Takuya Akiyama, Paulo Leal, Ceci Li, Leanne Well and members of the Microscopy Core for technical support, and the members of the Workman, Hawley and Mohan Laboratories for their input, discussion and feedback on the manuscript. This work was supported by funding from the Stowers Institute for Medical Research and a grant from the National Institute of General Medical Sciences (grant no. R35GM118068) to JLW.

## Supplemental information

**S1 Table. List of genotypes**. This table includes an overview of the Drosophila stocks that were used in this study.

**S2 Table. List of primers**. This table includes the sequences of the PCR primers used in this study.

**S3 Table. Motif Sequences**. Promoter groups were defined based on presence or absence of the different motifs. Each motif was scanned within the start and end windows using the TSS as a reference. A positive match for either DPE_O [71] or DPE_K [72] was considered a positive hit for a DPE element, as these are reported variants of the same motif.

**Fig S1.**
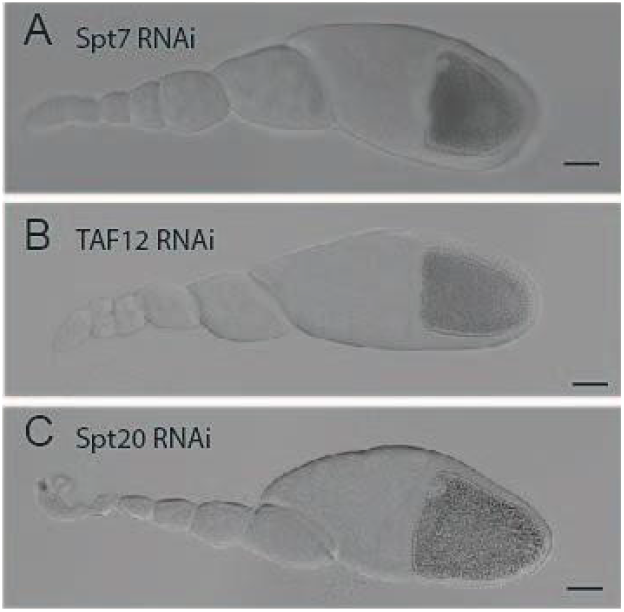
Different subunits of the SAGA core module are required in the germ line during oogenesis. Differential contrast images of ovarioles where SAGA core subunits have been depleted specifically in the germ line by UAS-Gal4 induced expression of RNAi constructs. C: RNAi Spt7 ovariole (29°C). D: RNAi TAF12 ovariole (29°C). E: RNAi Spt20 ovarioles (29°C). F: OreR control ovariole. Scale bar: 50 uM.

**Fig S2.**
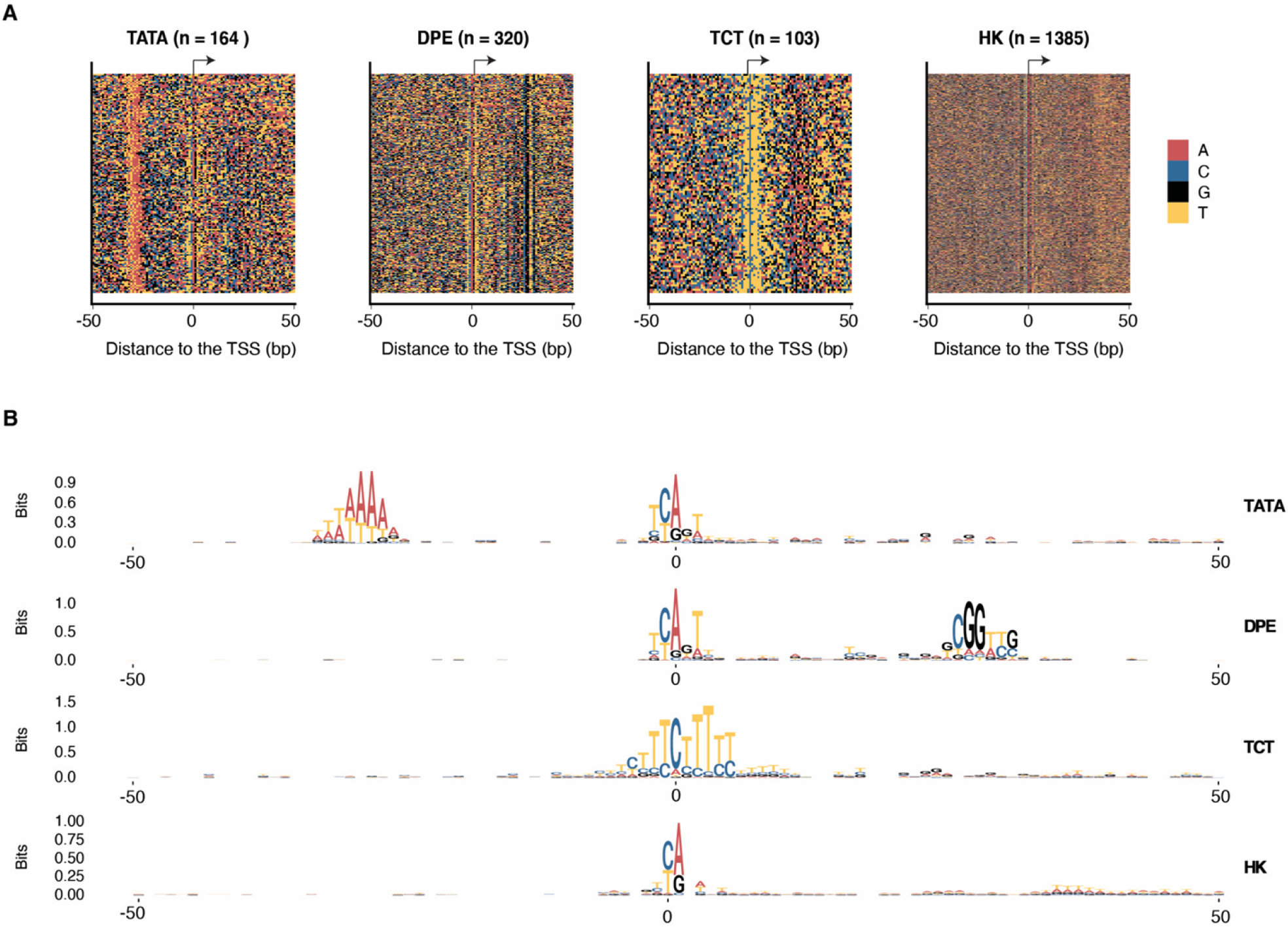
Define KC167 cell genes by promoter type. The DNA sequence of each promoter type is shown on the left as colored letters for a 101 bp window centered around the transcription start site. The position weight matrix (PWM) logo for each promoter type is displayed on the right.

**Fig S3.**
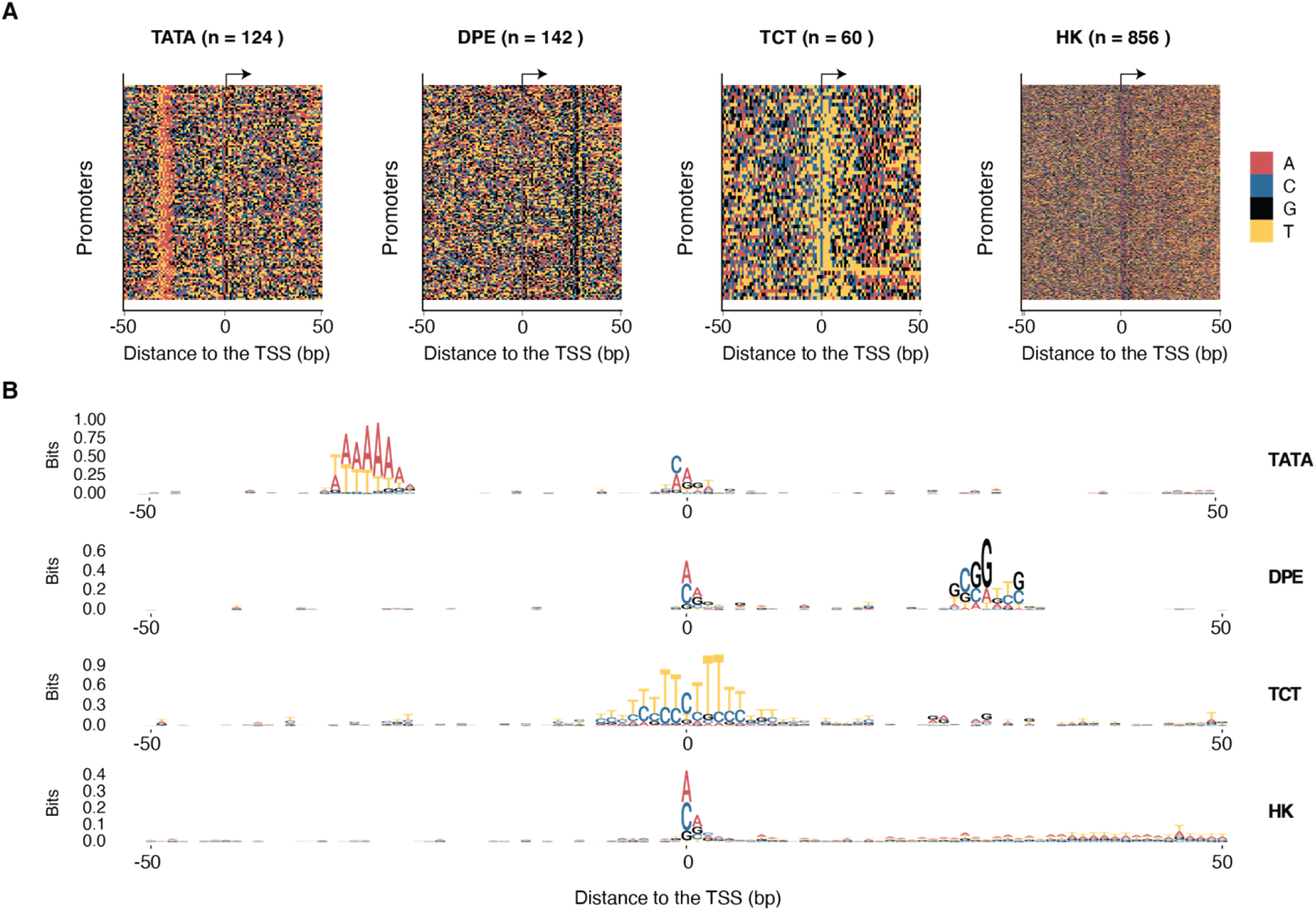
Define ovary cell genes by promoter type. The sequence of each promoter type is shown on the left as colored letters for a 101 bp window centered on the transcription start site. The consensus motif of each promoter type is displayed as logo on the right.

**Fig S4.**
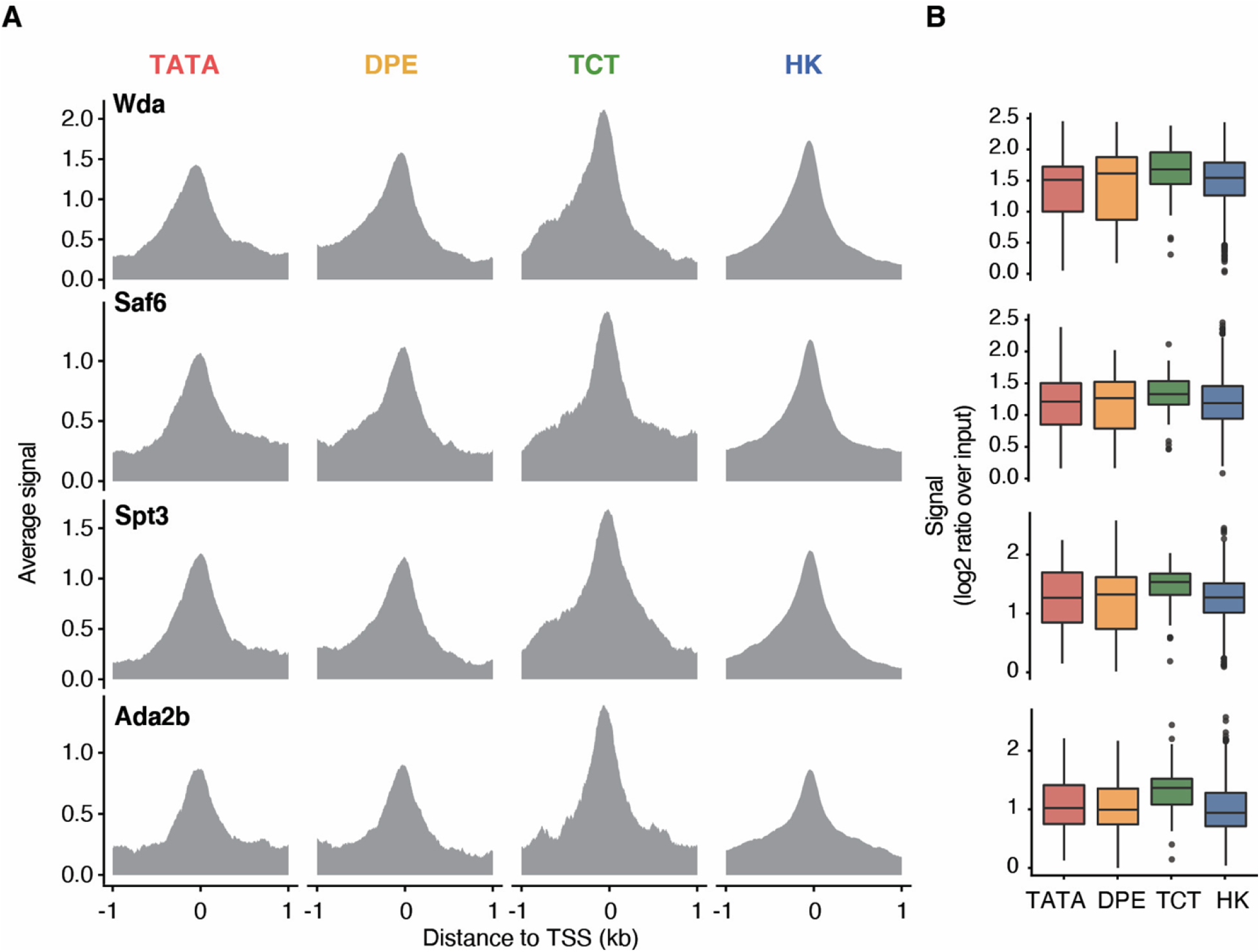
SAGA binding pattern at different promoter types in ovary tissue. (A) Average footprints of SAGA subunits in reads per million (RPM) for clusters of genes with different promoter types (left to right TATA, DPE-like, TCT and HK genes) per SAGA complex subunit (top to bottom SAGA subunits Ada2b, Spt3, SAF6 and WDA). (B) Boxplot displaying total signal per promoter type.

## References

1. Baptista T, Grunberg S, Minoungou N, Koster MJE, Timmers HTM, Hahn S, et al. SAGA Is a General Cofactor for RNA Polymerase II Transcription. Mol Cell. 2017.

2. Bian C, Xu C, Ruan J, Lee KK, Burke TL, Tempel W, et al. Sgf29 binds histone H3K4me2/3 and is required for SAGA complex recruitment and histone H3 acetylation. EMBO J. 2011;30(14):2829–42.

3. Grant PA, Duggan L, Cote J, Roberts SM, Brownell JE, Candau R, et al. Yeast Gcn5 functions in two multisubunit complexes to acetylate nucleosomal histones: characterization of an Ada complex and the SAGA (Spt/Ada) complex. Genes Dev. 1997;11(13):1640–50.

4. Candau R, Zhou JX, Allis CD, Berger SL. Histone acetyltransferase activity and interaction with ADA2 are critical for GCN5 function in vivo. EMBO J. 1997;16(3):555–65.

5. Grant PA, Eberharter A, John S, Cook RG, Turner BM, Workman JL. Expanded lysine acetylation specificity of Gcn5 in native complexes. J Biol Chem. 1999;274(9):5895–900.

6. Daniel JA, Torok MS, Sun ZW, Schieltz D, Allis CD, Yates JR, 3rd, et al. Deubiquitination of histone H2B by a yeast acetyltransferase complex regulates transcription. J Biol Chem. 2004;279(3):1867–71.

7. Serrano-Quílez J, Roig-Soucase S, Rodríguez-Navarro S. Sharing Marks: H3K4 Methylation and H2B Ubiquitination as Features of Meiotic Recombination and Transcription. Int J Mol Sci. 2020;21(12).

8. Brown CE, Howe L, Sousa K, Alley SC, Carrozza MJ, Tan S, et al. Recruitment of HAT complexes by direct activator interactions with the ATM-related Tra1 subunit. Science. 2001;292(5525):2333–7.

9. Han Y, Luo J, Ranish J, Hahn S. Architecture of the Saccharomyces cerevisiae SAGA transcription coactivator complex. EMBO J. 2014;33(21):2534–46.

10. Li H, Cuenin C, Murr R, Wang ZQ, Herceg Z. HAT cofactor Trrap regulates the mitotic checkpoint by modulation of Mad1 and Mad2 expression. Embo j. 2004;23(24):4824–34.

11. Park J, Kunjibettu S, McMahon SB, Cole MD. The ATM-related domain of TRRAP is required for histone acetyltransferase recruitment and Myc-dependent oncogenesis. Genes Dev. 2001;15(13):1619–24.

12. Fishburn J, Mohibullah N, Hahn S. Function of a eukaryotic transcription activator during the transcription cycle. Mol Cell. 2005;18(3):369–78.

13. Reeves WM, Hahn S. Targets of the Gal4 transcription activator in functional transcription complexes. Mol Cell Biol. 2005;25(20):9092–102.

14. Bhaumik SR, Green MR. SAGA is an essential in vivo target of the yeast acidic activator Gal4p. Genes Dev. 2001;15(15):1935–45.

15. Murr R, Vaissière T, Sawan C, Shukla V, Herceg Z. Orchestration of chromatin-based processes: mind the TRRAP. Oncogene. 2007;26(37):5358–72.

16. Lin L, Chamberlain L, Zhu LJ, Green MR. Analysis of Gal4-directed transcription activation using Tra1 mutants selectively defective for interaction with Gal4. Proc Natl Acad Sci U S A. 2012;109(6):1997–2002.

17. Helmlinger D, Marguerat S, Villén J, Swaney DL, Gygi SP, Bähler J, et al. Tra1 has specific regulatory roles, rather than global functions, within the SAGA co-activator complex. Embo j. 2011;30(14):2843–52.

18. Timmers HTM. SAGA and TFIID: Friends of TBP drifting apart. Biochimica et biophysica acta Gene regulatory mechanisms. 2020:194604.

19. Donczew R, Warfield L, Pacheco D, Erijman A, Hahn S. Two roles for the yeast transcription coactivator SAGA and a set of genes redundantly regulated by TFIID and SAGA. Elife. 2020;9.

20. Lee TI, Causton HC, Holstege FC, Shen WC, Hannett N, Jennings EG, et al. Redundant roles for the TFIID and SAGA complexes in global transcription. Nature. 2000;405(6787):701–4.

21. Papai G, Frechard A, Kolesnikova O, Crucifix C, Schultz P, Ben-Shem A. Atomic structure of the SAGA complex and it’s interaction with TBP. C R Biol. 2021;343(3):247–55.

22. Wang H, Dienemann C, Stutzer A, Urlaub H, Cheung ACM, Cramer P. Structure of the transcription coactivator SAGA. Nature. 2020.

23. Herbst DA, Esbin MN, Louder RK, Dugast-Darzacq C, Dailey GM, Fang Q, et al. Structure of the human SAGA coactivator complex: The divergent architecture of human SAGA allows modular coordination of transcription activation and co-transcriptional splicing. bioRxiv. 2021:2021.02.08.430339.

24. Carre C, Szymczak D, Pidoux J, Antoniewski C. The histone H3 acetylase dGcn5 is a key player in Drosophila melanogaster metamorphosis. Mol Cell Biol. 2005;25(18):8228–38.

25. Li X, Seidel CW, Szerszen LT, Lange JJ, Workman JL, Abmayr SM. Enzymatic modules of the SAGA chromatin-modifying complex play distinct roles in Drosophila gene expression and development. Genes Dev. 2017;31(15):1588–600.

26. Soffers JHM, Li X, Saraf A, Seidel CW, Florens L, Washburn MP, et al. Characterization of a metazoan ADA acetyltransferase complex. Nucleic Acids Res. 2019.

27. Torres-Zelada EF, Stephenson RE, Alpsoy A, Anderson BD, Swanson SK, Florens L, et al. The Drosophila Dbf4 ortholog Chiffon forms a complex with Gcn5 that is necessary for histone acetylation and viability. J Cell Sci. 2018.

28. Rabenstein MD, Zhou S, Lis JT, Tjian R. TATA box-binding protein (TBP)-related factor 2 (TRF2), a third member of the TBP family. Proc Natl Acad Sci U S A. 1999;96(9):4791–6.

29. Nakamura S, Hira S, Kojima M, Kondo A, Mukai M. Expression of the core promoter factors TATA box binding protein and TATA box binding protein-related factor 2 in Drosophila germ cells and their distinct functions in germline development. Dev Growth Differ. 2020;62(9):540–53.

30. Kopytova DV, Krasnov AN, Kopantceva MR, Nabirochkina EN, Nikolenko JV, Maksimenko O, et al. Two isoforms of Drosophila TRF2 are involved in embryonic development, premeiotic chromatin condensation, and proper differentiation of germ cells of both sexes. Mol Cell Biol. 2006;26(20):7492–505.

31. Martianov I, Velt A, Davidson G, Choukrallah MA, Davidson I. TRF2 is recruited to the pre-initiation complex as a testis-specific subunit of TFIIA/ALF to promote haploid cell gene expression. Sci Rep. 2016;6:32069.

32. Metcalf CE, Wassarman DA. Nucleolar colocalization of TAF1 and testis-specific TAFs during Drosophila spermatogenesis. Developmental dynamics : an official publication of the American Association of Anatomists. 2007;236(10):2836–43.

33. Pointud JC, Mengus G, Brancorsini S, Monaco L, Parvinen M, Sassone-Corsi P, et al. The intracellular localisation of TAF7L, a paralogue of transcription factor TFIID subunit TAF7, is developmentally regulated during male germ-cell differentiation. J Cell Sci. 2003;116(Pt 9):1847–58.

34. Zhou H, Grubisic I, Zheng K, He Y, Wang PJ, Kaplan T, et al. Taf7l cooperates with Trf2 to regulate spermiogenesis. Proc Natl Acad Sci U S A. 2013;110(42):16886–91.

35. Hiller M, Chen X, Pringle MJ, Suchorolski M, Sancak Y, Viswanathan S, et al. Testis-specific TAF homologs collaborate to control a tissue-specific transcription program. Development. 2004;131(21):5297–308.

36. Gustafson EA, Seymour KA, Sigrist K, Rooij D, Freiman RN. ZFP628 Is a TAF4b-Interacting Transcription Factor Required for Mouse Spermiogenesis. Mol Cell Biol. 2020;40(7).

37. Langer D, Martianov I, Alpern D, Rhinn M, Keime C, Dollé P, et al. Essential role of the TFIID subunit TAF4 in murine embryogenesis and embryonic stem cell differentiation. Nat Commun. 2016;7:11063.

38. Grive KJ, Gustafson EA, Seymour KA, Baddoo M, Schorl C, Golnoski K, et al. TAF4b Regulates Oocyte-Specific Genes Essential for Meiosis. PLoS Genet. 2016;12(6):e1006128.

39. Yu C, Cvetesic N, Hisler V, Gupta K, Ye T, Gazdag E, et al. TBPL2/TFIIA complex establishes the maternal transcriptome through oocyte-specific promoter usage. Nat Commun. 2020;11(1):6439.

40. Weake VM, Swanson SK, Mushegian A, Florens L, Washburn MP, Abmayr SM, et al. A novel histone fold domain-containing protein that replaces TAF6 in Drosophila SAGA is required for SAGA-dependent gene expression. Genes Dev. 2009;23(24):2818–23.

41. Georgieva S, Kirschner DB, Jagla T, Nabirochkina E, Hanke S, Schenkel H, et al. Two novel Drosophila TAF(II)s have homology with human TAF(II)30 and are differentially regulated during development. Mol Cell Biol. 2000;20(5):1639–48.

42. Guelman S, Suganuma T, Florens L, Weake V, Swanson SK, Washburn MP, et al. The essential gene wda encodes a WD40 repeat subunit of Drosophila SAGA required for histone H3 acetylation. Mol Cell Biol. 2006;26(19):7178–89.

43. He Q, Johnston J, Zeitlinger J. ChIP-nexus enables improved detection of in vivo transcription factor binding footprints. Nat Biotechnol. 2015;33(4):395–401.

44. Bastock R, St Johnston D. Drosophila oogenesis. Curr Biol. 2008;18(23):R1082–7.

45. Papai G, Frechard A, Kolesnikova O, Crucifix C, Schultz P, Ben-Shem A. Structure of SAGA and mechanism of TBP deposition on gene promoters. Nature. 2020.

46. Wragg JW, Roos L, Vucenovic D, Cvetesic N, Lenhard B, Müller F. Embryonic tissue differentiation is characterized by transitions in cell cycle dynamic-associated core promoter regulation. Nucleic Acids Res. 2020;48(15):8374–92.

47. Lu D, Sin HS, Lu C, Fuller MT. Developmental regulation of cell type-specific transcription by novel promoter-proximal sequence elements. Genes Dev. 2020;34(9-10):663–77.

48. Vo Ngoc L, Wang YL, Kassavetis GA, Kadonaga JT. The punctilious RNA polymerase II core promoter. Genes Dev. 2017;31(13):1289–301.

49. Chen K, Johnston J, Shao W, Meier S, Staber C, Zeitlinger J. A global change in RNA polymerase II pausing during the Drosophila midblastula transition. Elife. 2013;2:e00861.

50. de Jonge WJ, O’Duibhir E, Lijnzaad P, van Leenen D, Groot Koerkamp MJ, Kemmeren P, et al. Molecular mechanisms that distinguish TFIID housekeeping from regulatable SAGA promoters. Embo j. 2017;36(3):274–90.

51. Tora L, Timmers HT. The TATA box regulates TATA-binding protein (TBP) dynamics in vivo. Trends Biochem Sci. 2010;35(6):309–14.

52. Celniker SE, Dillon LA, Gerstein MB, Gunsalus KC, Henikoff S, Karpen GH, et al. Unlocking the secrets of the genome. Nature. 2009;459(7249):927–30.

53. Kedmi A, Zehavi Y, Glick Y, Orenstein Y, Ideses D, Wachtel C, et al. Drosophila TRF2 is a preferential core promoter regulator. Genes Dev. 2014;28(19):2163–74.

54. Vo Ngoc L, Kassavetis GA, Kadonaga JT. The RNA Polymerase II Core Promoter in Drosophila. Genetics. 2019;212(1):13–24.

55. Zabidi MA, Arnold CD, Schernhuber K, Pagani M, Rath M, Frank O, et al. Enhancer-core-promoter specificity separates developmental and housekeeping gene regulation. Nature. 2015;518(7540):556–9.

56. Wang YL, Duttke SH, Chen K, Johnston J, Kassavetis GA, Zeitlinger J, et al. TRF2, but not TBP, mediates the transcription of ribosomal protein genes. Genes Dev. 2014;28(14):1550–5.

57. Moore PA, Ozer J, Salunek M, Jan G, Zerby D, Campbell S, et al. A human TATA binding protein-related protein with altered DNA binding specificity inhibits transcription from multiple promoters and activators. Mol Cell Biol. 1999;19(11):7610–20.

58. Teichmann M, Wang Z, Martinez E, Tjernberg A, Zhang D, Vollmer F, et al. Human TATA-binding protein-related factor-2 (hTRF2) stably associates with hTFIIA in HeLa cells. Proc Natl Acad Sci U S A. 1999;96(24):13720–5.

59. Hochheimer A, Zhou S, Zheng S, Holmes MC, Tjian R. TRF2 associates with DREF and directs promoter-selective gene expression in Drosophila. Nature. 2002;420(6914):439–45.

60. Shao W, Zeitlinger J. Paused RNA polymerase II inhibits new transcriptional initiation. Nat Genet. 2017;49(7):1045–51.

61. Bonnet J, Wang CY, Baptista T, Vincent SD, Hsiao WC, Stierle M, et al. The SAGA coactivator complex acts on the whole transcribed genome and is required for RNA polymerase II transcription. Genes Dev. 2014;28(18):1999–2012.

62. Dahlberg O, Shilkova O, Tang M, Holmqvist PH, Mannervik M. P-TEFb, the super elongation complex and mediator regulate a subset of non-paused genes during early Drosophila embryo development. PLoS Genet. 2015;11(2):e1004971.

63. Saint M, Sawhney S, Sinha I, Singh RP, Dahiya R, Thakur A, et al. The TAF9 C-terminal conserved region domain is required for SAGA and TFIID promoter occupancy to promote transcriptional activation. Mol Cell Biol. 2014;34(9):1547–63.

64. Weake VM, Dyer JO, Seidel C, Box A, Swanson SK, Peak A, et al. Post-transcription initiation function of the ubiquitous SAGA complex in tissue-specific gene activation. Genes Dev. 2011;25(14):1499–509.

65. Hassan AH, Prochasson P, Neely KE, Galasinski SC, Chandy M, Carrozza MJ, et al. Function and selectivity of bromodomains in anchoring chromatin-modifying complexes to promoter nucleosomes. Cell. 2002;111(3):369–79.

66. Ringel AE, Cieniewicz AM, Taverna SD, Wolberger C. Nucleosome competition reveals processive acetylation by the SAGA HAT module. Proc Natl Acad Sci U S A. 2015;112(40):E5461–70.

67. Kondo S, Ueda R. Highly improved gene targeting by germline-specific Cas9 expression in Drosophila. Genetics. 2013;195(3):715–21.

68. Huang F, Paulson A, Dutta A, Venkatesh S, Smolle M, Abmayr SM, et al. Histone acetyltransferase Enok regulates oocyte polarization by promoting expression of the actin nucleation factor spire. Genes Dev. 2014;28(24):2750–63.

69. Haberle V, Forrest AR, Hayashizaki Y, Carninci P, Lenhard B. CAGEr: precise TSS data retrieval and high-resolution promoterome mining for integrative analyses. Nucleic Acids Res. 2015;43(8):e51.

70. Edgar R, Domrachev M, Lash AE. Gene Expression Omnibus: NCBI gene expression and hybridization array data repository. Nucleic Acids Res. 2002;30(1):207–10.

71. Ohler U, Liao GC, Niemann H, Rubin GM. Computational analysis of core promoters in the Drosophila genome. Genome Biol. 2002;3(12):Research0087.

72. Burke TW, Kadonaga JT. Drosophila TFIID binds to a conserved downstream basal promoter element that is present in many TATA-box-deficient promoters. Genes Dev. 1996;10(6):711–24.

